# Efficient RNA Folding Simulation via a Structure-Based Single-Site-Per-Nucleotide Model

**DOI:** 10.64898/2025.12.13.694107

**Authors:** Thomas Thornton, Xingcheng Lin

## Abstract

Computational modeling of large RNA structures and their dynamics is essential for uncovering the molecular mechanisms underlying various genomic processes and RNA-regulated cellular functions. Residue-resolution modeling is an effective approach for simulating large biomolecular structures while preserving essential sequence and structural features presented in atomic structures. Here, we implemented a structure-based single-site-per-nucleotide (SSPN) RNA model using the GPU-accelerated OpenMM ^1^ software and evaluated its computational efficiency and accuracy by simulating RNA hairpins unfolding under force. Our simulations compare favorably with an earlier, more detailed RNA model and quantitatively reproduce experimentally measured thermodynamic properties of RNA under mechanical stretching. This SSPN model enables scalable and accurate simulations of large RNA ensembles, such as long non-coding RNAs and RNA liquid-liquid phase separation.

## Introduction

RNA is a single-stranded nucleic acid that constitutes the majority of the human genome,^2^ and its structures play critical roles in regulating various cellular functions, from transcription and translation,^3–5^ to gene expression^6–8^ and the formation of RNA-based biomolecular condensates.^9,10^ RNA adopts heterogeneous secondary and tertiary structures under varying cellular conditions,^11^ and these conformations are strongly influenced by the surrounding environment and interactions with other cellular components.

Computational modeling and simulations have become important tools for elucidating RNA dynamics and their functional consequences.^12–15^ Significant progress has been made in both atomistic and coarse-grained models for characterizing RNA folding^14–22^ and its modulation by environmental factors, such as metal-ion binding.^23–25^ Among those developments, the structure-based model^26–28^ – grounded in the principle of minimal frustration^29,30^ – enables efficient samplings of biomolecular dynamics by assuming an unfrustrated funneled folding energy landscape.^31,32^ It has been successfully applied to studying protein folding,^33–35^ protein structural transition,^36–39^ and the prediction of sequence-specific mutational effects.^26^ More recently, structure-based models have been extended to RNA, elucidating its folding dynamics ^12,14,17,40^ and its modulation by metal ions,^16,25,41^ offering mechanistic insights into RNA structural dynamics.

Structure-based RNA models have been implemented at multiple resolutions, including all-atom resolution,^12,14,16^ three-site-per-nucleotide models,^17,40^ and single-site-per-nucleotide (SSPN) models.^42,43^ Among these, an SSPN model can most efficiently simulate large RNA structures, while retaining essential polymeric features and backbone dynamics. A residue-level resolution may preserve both the structural and sequence identities of RNA, enabling accurate modeling of biomolecular surfaces and interactions with other cellular components, such as proteins ^44^ and ions.^24^

In this manuscript, we implemented an SSPN model using the GPU-accelerated OpenMM computational platform^1^ and benchmarked its computational efficiency across multiple CPU and GPU cores. We further evaluate the model’s accuracy by comparing its predictions to those from a more detailed three-site-per-nucleotide RNA simulation model and to single-molecule force spectroscopy by revisiting the folding and unfolding of *Tetrahymena thermophila ribozyme*.^17,40,42,45,46^ Our results demonstrate that the SSPN model can reproduce the essential RNA folding thermodynamics captured by a detailed simulation and provide additional structural insights into folding pathways underlying experimental observations. Together, our studies demonstrate that a GPU-implemented SSPN RNA model is capable of simulating large RNA bimolecules, including typical long non-coding RNAs (lncRNAs)^6,7^ and RNAs involved in liquid-liquid phase separation.^10,47–49^

## Materials and Methods

### Structure-based single-site-per-nucleotide RNA model

We implemented the unfrustrated, structure-based single-site-per-nucleotide (SSPN) RNA model^13,29,30^ based on the GPU-accelerated OpenMM simulation framework.^1^ This GPU-accelerated model achieves high computational efficiency while preserving RNA topology. It employs the structure-based SMOG force field,^28^ adapted by modifying modules of the OpenABC package.^50^ Specifically, each nucleotide is represented by a single coarse-grained (CG) site located at its phosphorus atom. This CG representation is consistent with the single-site CG protein model, which represents each amino acid by a single site at the C*_α_* atom position.

Short-range bonded interactions were used to preserve the polymeric physics of RNA, while native interactions – adopted from the original protein force field – were used to maintain the structural integrity and thermodynamic stability of the molecules. As a first-order approximation, we hypothesize that the topology of the native structure, rather than sequence-specific energetic heterogeneity, determines RNA folding properties in this unfrustrated model. Accordingly, all native nucleotide pairs are assigned a uniform interaction strength. A similar approach has been successfully implemented in prior protein folding models.^26–28^

Following earlier studies that used structure-based models in proteins and protein-DNA systems,^14,26–28,51–55^ the total potential energy for an N-nucleotide RNA chain is expressed as

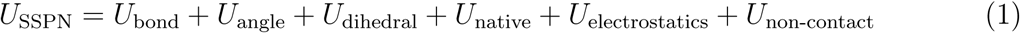

The bond, angle, and dihedral terms preserve chain connectivity and are minimized at their native values.

The bond term, *U*_bond_ is harmonic, and can be expressed as

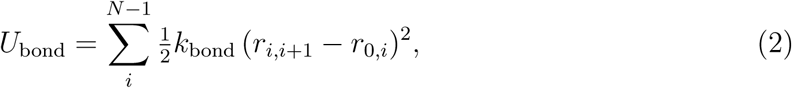

where *r*_0*,i*_ are the native bond length, and the force constant *k*_bond_= 75,000 kJ/(mol·nm^2^). The angle term, *U*_angle_, is also harmonic. For each bead, it acts on two neighboring beads

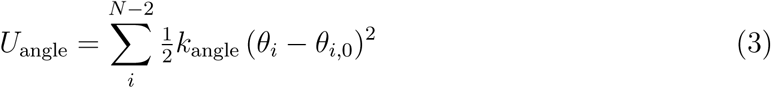

where *θ_i_* is the angle, *θ_i,_*_0_ is the native angle, and the force constant *k*_angle_ = 150 kJ/mol/rad^2^. To improve the system stability, we implemented an angle clipping feature described below.

### The dihedral term, *U*_dihedral_, captures torsion and includes two periodic components with periodicity 1 and 3, respectively

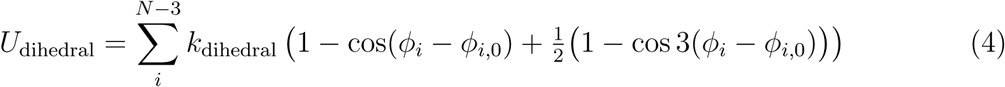

where *ϕ_i_* is the dihedral angle (the angles between the two planes which pass through two sets of three coarse-grained sites, having two sites in common; see Fig. 1), *ϕ_i,_*_0_ is the native dihedral angle, and *k*_dihedral_ = 3.75 kJ*/*(mol · rad^2^).

**Figure 1:**
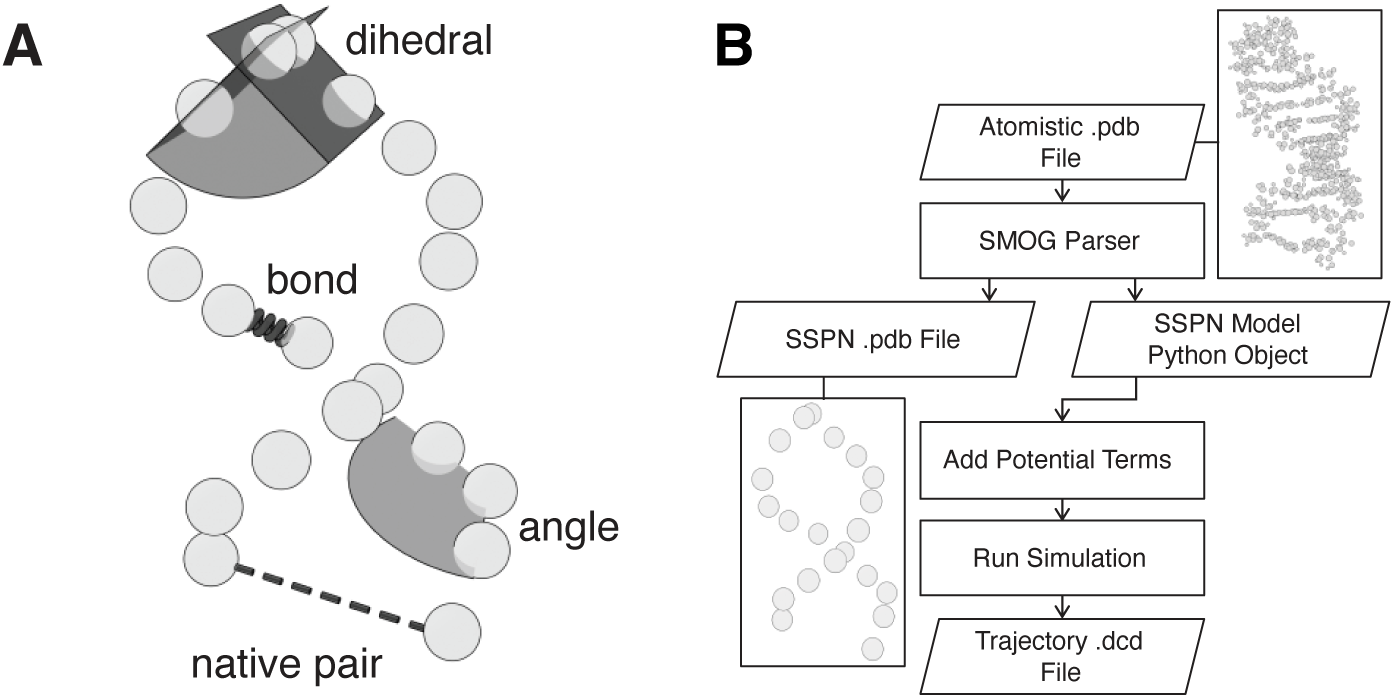
Schematic illustration and workflow of the SSPN model. (A) Coarse-grained representation of the SSPN model, illustrating the definition of bond, angle, dihedral, and native pairs. (B) Workflow of the SSPN model, which takes as input an atomistic RNA PDB structure, converts it into a P-atom-based coarse-grained representation, and performs simulations using a structure-based potential.

The native pairing term, *U*_native_, is a long-range interaction that stabilizes helices as well as secondary and tertiary structures. It occurs between pairs of residues that are spatially close in the native structure but do not interact via the chain connectivity terms. Inspired by an earlier approach in modeling structure-based protein model,^56^ we model the native interaction using a Gaussian potential centered on the pair’s equilibrium separation,

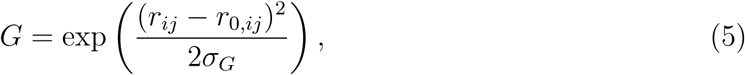

where *σ_G_* = 0.05. The potential itself has the mathematical form:

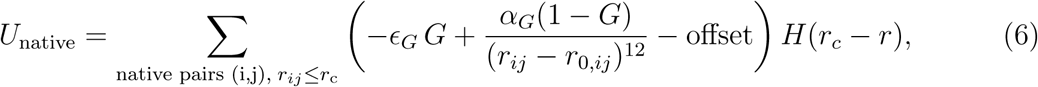

where *r_ij_* is the separation between the two beads, *r*_0*,ij*_ is the their equilibrium separation, *ɛ_G_* = 3.75 kJ/mol/nm^2^, *α_G_* = 6.29 × 10*^−^*^5^ kJ/mol/nm^2^. *H* represents the Heaviside step function. The cutoff distance *r*_c_ = 6*σ_G_* · *r*_0*,ij*_, and offset = −*ɛ_G_*

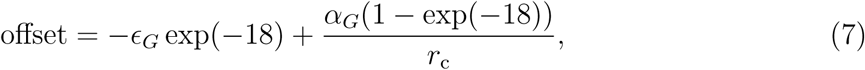

which ensures continuity at the cutoff distance.

The electrostatic term, *U*_electrostatics_, is modeled by a Debye-Hückel potential times a switching function

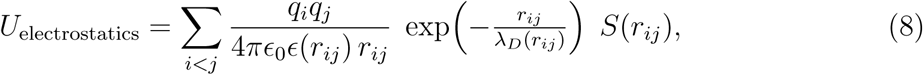

The Debye-Hückel potential takes into account the effects of ion shielding, and uses a distance-dependent dielectric constant,

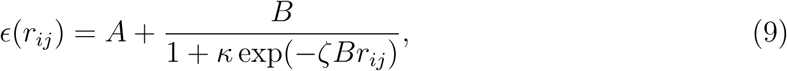

where *A* = −8.5525, *κ* = 7.7839, *B* = *ɛ*_water_ − *A*, and *ζ* = 0.0362 nm*^−^*^1^. These values allow the dielectric constant to be that of water at large distances and that found inside bulk protein at short distances. The Debye length *λ_D_*(*r_ij_*) determines the rate of falloff due to ion screening,

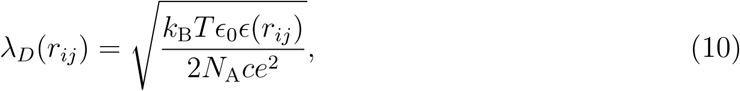

where *c* is the salt concentration used in the simulation and was set to be 150 mM in all the studies reported in this work. The switching function *S*(*r_ij_*) is a fifth-order spline, where the potential remains unaffected at distances less than 1.2 nm and decays smoothly, reaching zero at *r_ij_* = 1.5 nm.

The non-contact term, *U*_NC_, represents the excluded volume potential and is modeled with the repulsive part of the Lennard-Jones potential

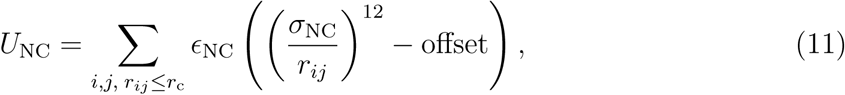

#### Structure-based angle clipping

To ensure the stability of molecules during simulation, we incorporated the angle-clipping feature from the maximum entropy optimized force field (MOFF) model.^57^ This puts a ceiling on the allowed equilibrium angles taken from the native structure. All equilibrium angles exceeding 160°, an empirical number, are clipped to this value.

#### Determination of native pairs

To define the native pairing interaction, a contact map was determined based on the Shadow algorithm^58^ implemented in the OpenABC framework.^50^ First, contacts between heavy atoms were identified from the atomistic structure. A CG native nucleotide pair was defined when the minimum atomic separation between two nucleotides was smaller than 6°A. All CG native interactions were assigned the same interaction strength.

### Implementation of SSPN in OpenMM

The SSPN RNA model was implemented by modifying the OpenABC package, ^50^ which is based on the OpenMM computational platform.^1^ Figure 1 illustrates the workflow. To set up the simulation, the all-atom PDB is parsed to generate a CG PDB containing only the phosphorus atoms from the atomistic PDB. During parsing, the equilibrium interaction parameters of the force field’s interaction terms are directly extracted from the structure. The SSPN RNA can then be copied and inserted randomly (without overlaps) into the simulation box multiple times. For all simulations in this study, a single RNA molecule was inserted, except for the efficiency benchmarking (details in Section *Benchmark model efficiency*). Finally, one can add interaction terms based on the inserted molecules before launching the simulation. Example scripts and detailed instructions are available at the GitHub repository https://github.com/LinResearchGroup-NCSU/SSPN_RNA_Model.

### Simulation protocol

All simulations were performed using Langevin dynamics at 300 K with a timestep of 10 fs and a friction coefficient of 1.0 ps*^−^*^1^. The 10 fs timestep was used as it maintains the system’s stability (Figure S1). Ionic concentrations were implicitly simulated with the Debye-Hückel treatment of screening effects, and salt concentration is set to be 150 mM in all simulations. Except for the benchmarking simulations, all simulations reported in this study were run on a single NVIDIA L40s GPU.

#### Benchmark model efficiency

To evaluate computational efficiency, we simulated systems containing 100, 500, 1000, and 5000 copies of the 76-nt P5abc RNA (extracted from the *Tetrahymena thermophila ribozyme*, modeling details are provided in Section *Modeling and simulations of P5ab, P5abc*ΔA*, and P5abc systems.*). Simulations were run on Intel Xeon Platinum 8462Y+ CPUs and on a single NVIDIA L40s GPU. Performance was measured as mean simulated nanoseconds per day, averaged over the latter half of 0.4 ns trajectories (40,000 steps) to ensure the systems were fully equilibrated.

#### Modeling and simulations of the P5GA hairpin

We performed constant force simulations of the 22-nucleotide P5GA hairpin (PDB ID: 1F9L), which represents a simplified version of the P5abc region of a Group I Ribozyme.^17,59^ Because the experimental structure lacks the 5’-terminal phosphate, our simulation contained 21 of the 22 nucleotides. For constant force simulations, we applied the OpenMM command `CustomCentroidBondForce` between the first and last residues of the RNA.

To compute the equilibrium free energy profiles at different pulling forces, we applied forces ranging from 0 pN to 8 pN for one 1 µs, with data collected every 200 steps. To calculate the equilibrium constant *K*_eq_(*f*), we took a more dense sampling of forces near the unfolding force, from 2.5 pN to 7.0 pN in increments of 0.25 pN. Each simulation lasts 100 ns with a reporting interval of 5000 time steps.

#### Modeling and simulations of P5ab, P5abcΔA, and P5abc systems

In addition to the P5GA hairpin, we modeled and simulated the P5abc domain of the *Tetrahymena thermophila* group I intron, as well as its two truncated variants. The all-atom P5abc structure was generated by trimming the full P4-P6 domain of the *Tetrahymena thermophila ribozyme* (PDB ID: 1GID^60^). We then created all-atom PDB files for the two truncated variants – P5abcΔA and P5ab – that were examined experimentally in Reference 45. P5abcΔA lacks the A-rich bulge present in P5abc, and P5ab lacks the entire P5c helix, making it a hairpin. Since experimental structures for these variations were not available, we approximated them by modifying the atomistic P5abc PDB: We first edited the PDB file to remove the appropriate residues to match the sequences reported in Figure 1 of Reference 45. The resulting discontinuities in the RNA strand were repaired by manually repositioning the terminal residues using VMD (Visual Molecular Dynamics) software.^61^ We did not alter the angular orientation of the repositioned residues, as such changes have a negligible effect on the CG structure.^∗^ Finally, residue indices were renumbered to restore continuous indexing in the final PDB files.

To compute the unfolding free energy, Δ*G*, for all three systems, we performed umbrella sampling^62^ with the end-to-end distance as the biasing coordinate to gain enhanced sampling over the free-energy landscape. For each umbrella, a harmonic restraint was imposed between the first and last residues using PLUMED.^63^ Umbrellas were centered from 0 to 27 nm with an increment of 1 nm. For each simulation, the RNA was first stretched for 1.75 ns using a high force constant *k* = 50.0 kJ mol*^−^*^1^ nm^2^ to reach the umbrella center, after which the force constant was reduced to 5.0 kJ mol*^−^*^1^ nm^2^ and the simulation continued for another 40 ns in equilibrated state. When constructing the free energy profiles, we disregarded the initial 1.75 ns of simulation and used the Weighted Histogram Analysis Method (WHAM)^64^ implemented in the SMOG package.^28^

#### Constant velocity simulation of P5abc and P5abcΔA

To probe unfolding dynamics of the P5abc system as done in optical tweezer experiments, we performed constant-velocity pulling simulations. The optical trap was modeled as a harmonic restraint force between the first and last residues, 0.5*k*(*r* − *r*_0_)^2^, where *r*_0_ was moving with a fixed velocity *v*. To simulate the reversible folding/unfolding curves of P5abcΔA, the sign of the velocity becomes negative halfway through the simulation. We used the loading rate, *r* = *kv*, to control the rupture process, with *k* being the force constant of the optical trap. To estimate a physically realistic loading rate, we compared simulated and experimental hysteresis, defined as the area between the unfolding and folding branches of the force-extension curve, and quantified the extent to which the unfolding transition is irreversible under constant loading rate. Simulations were conducted across a range of loading rates, for 400 ns · 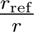. Hysteresis was computed from simulated force-extension trajectories and compared with experimental values extracted using WebPlotDigitizer.^65^ The reference loading rate *r*_ref_ was defined as the simulation loading rate that reproduced experimentally observed hysteresis. We then performed 200 independent constant-velocity pulling simulations of P5abc using *r*_ref_, each for 200 ns, to investigate unfolding dynamics.

## Results

### Efficient Structure-Based RNA Model at Single-Nucleotide Resolution

We implemented the single-site-per-nucleotide (SSPN) structure-based model using the GPU-optimized OpenMM platform^1^ implemented under the OpenABC framework. ^50^ Rooted in the principle of minimal frustration, ^29–31^ the model postulates an RNA’s folded structure as the energetic minimum of its funneled folding landscape. ^12,14,17,26^ By approximating solvent-averaged interactions between RNA molecules with stabilizing native contacts encoded in experimentally resolved structures, the SSPN model removes energetic frustration and provides a computationally efficient baseline model for RNA folding simulations (Figure 1A). Similar unfrustrated structure-based models have been successfully applied to protein folding,^26,27,56^ protein structural transitions,^36–39^ and RNA dynamics and function.^12,13,17,39^ Additional details are provided in the Materials and Methods Section *Structure-based single-site-per-nucleotide RNA model*.

The SSPN model uses Python scripting to simplify system construction (Figure 1B). The model takes as input an atomistic RNA folded structure, either downloaded from the PDB website ^66^ or predicted by deep-learning-based structure prediction tools.^67–70^ A Python parser, adapted from the SMOG scripts^28^ provided in OpenABC,^50^ was used to generate the structure-based model using the phosphorus atom of each nucleotide as the coarse-grained site. Both bonded and non-bonded native interactions, along with Debye-Hückel treatment of electrostatic interactions, were added as potential terms to construct the full potential function for simulations. The processed object was used for running simulations, with output trajectories in .dcd format. A detailed workflow is described in the Materials and Methods Section *Implementation of SSPN in OpenMM*.

### Efficient simulations with a GPU-based Computational Platform

To evaluate the computational efficiency of the SSPN model, we conducted comprehensive benchmarking across four independent systems with varying numbers of RNA molecules. Each system contains multiple copies of 76-residue P5abc RNA, (Figure 2). All simulations were run for 80,000 steps. Additional details are provided in Section *Benchmark model efficiency*.

**Figure 2:**
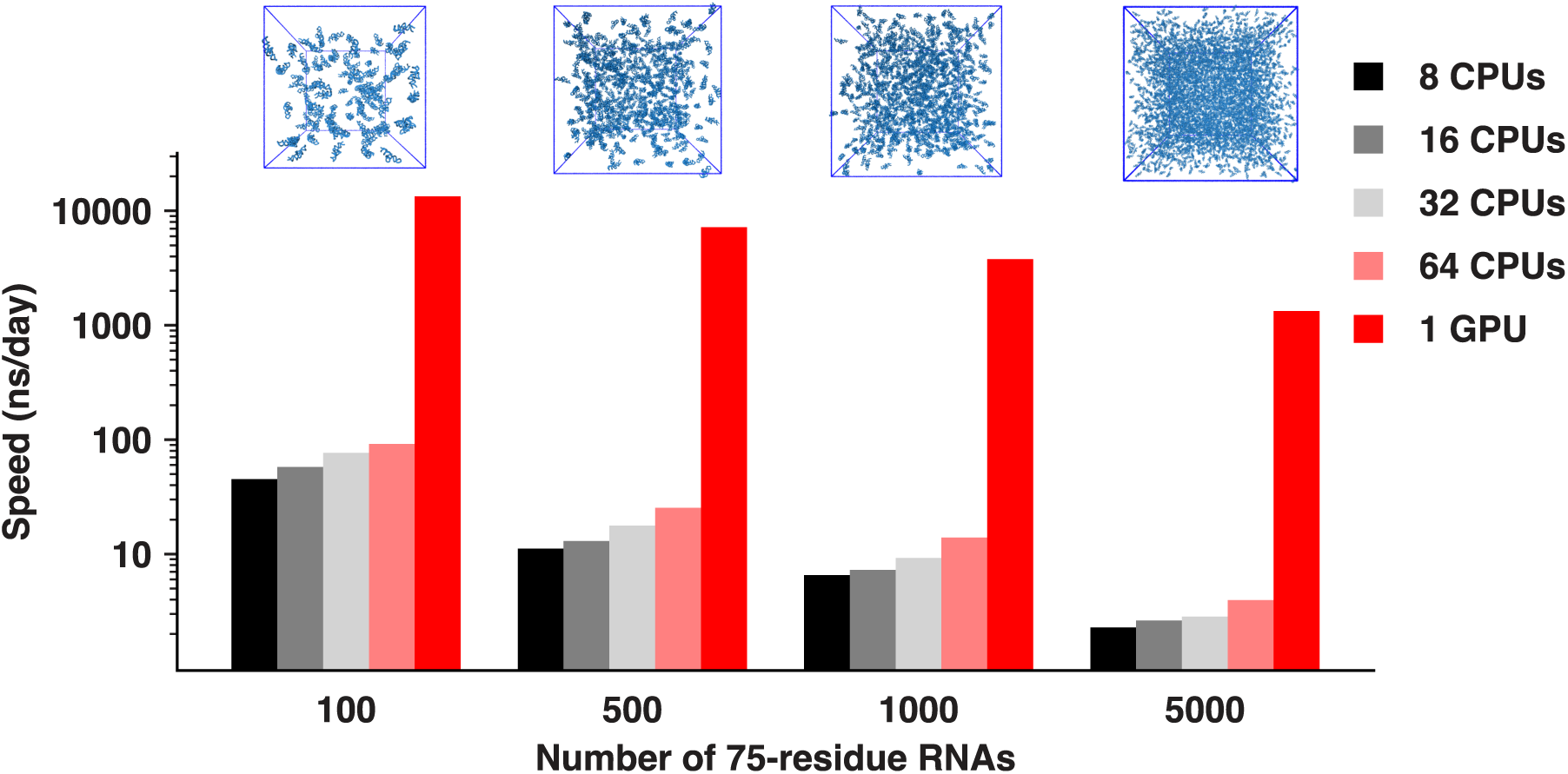
Computation speed. Snapshots of four systems containing 100-5000 copies of the 76-nt p5abc RNAs. Benchmark simulations were performed on Intel Xeon Platinum 8462Y+ CPUs and on a single NVIDIA L40s GPU. Bars represent computational speeds measured from simulations after equilibration.

As shown in Figure 2, GPU-based simulations significantly outperform those by CPU. To assess the scaling of simulation with increasing system size, we measured computational speed across systems with increasing RNA copy numbers. From 100 to 500 RNAs, CPU performance dropped to 25% of its initial value, whereas GPU performance dropped to 53% of its initial value. For larger systems (1000-5000 RNAs), CPU scaling improved to 32% of its initial value, while GPU scaling reduced to 35%. These results indicate that GPU is the optimal choice for systems up to 400k nucleotides, whereas CPUs may be more efficient for even larger numbers of nucleotides. Notably, even for the largest system tested (375k nucleotides), the SSPN model achieved over 1 *µs* of simulation time per day, demonstrating its efficiency in simulating large RNA systems, including those involved in liquid-liquid phase separation (LLPS).^10,47–49^

### SSPN recapitulates essential RNA folding properties from detailed RNA model

It is important to examine the SSPN model’s accuracy in characterizing key thermodynamic and kinetic features of RNA dynamics. To this end, we simulated the mechanical unfolding of P5GA (PDB ID: 1F9L), an RNA hairpin that contains the core sequence element of the P5abc region of the *Tetrahymena thermophila ribozyme*. The mechanical unfolding of P5GA has been extensively characterized using a detailed three-bead-per-nucleotide RNA model,^17^ which quantified both the thermodynamics and kinetics of P5GA folding and semi-quantitatively reproduces an earlier experimental measurement by optical tweezers^45^

To evaluate the accuracy of the SSPN model, we simulated the P5GA hairpin under a range of constant pulling forces. The model’s computational efficiency allows us to quantitatively capture the free energy of RNA hairpin unfolding from single-temperature simulations at 300K (Figure 3A). The results show a 3.3 *k*_B_*T* penalty in unfolding RNA and a decrease in this penalty as the applied force increases. We further calculated the equilibrium constant of RNA unfolding (Figure 3B) from the unfolding trajectories. The molecule is considered unfolded when its extension exceeds 5.0 nm, as observed from the free energy and trajectory plots at 4 pN, where folded and unfolded populations are similar (Figure S2). Fitting log(*K*_eq_) as a function applied forces gives log *K_eq_* = 0.62*f* − 2.82. Using the thermodynamic relation log *K*_eq_(*f*) = Δ*F*_UF_*/k*_B_*T* + *f* · Δ*x*_UF_*/k*_B_*T*, we obtain Δ*x*_UF_ = 2.55 nm and Δ*F*_UF_ = 2.82 *k*_B_*T*. The transition midpoint (*K*_eq_ = 1) gives *f* = 4.5 pN, consistent with the free energy profiles. Overall, the SSPN model quantitatively reproduces key thermodynamic features of RNA folding from the more detailed coarse-grained RNA model, despite using a simplified representation.

**Figure 3:**
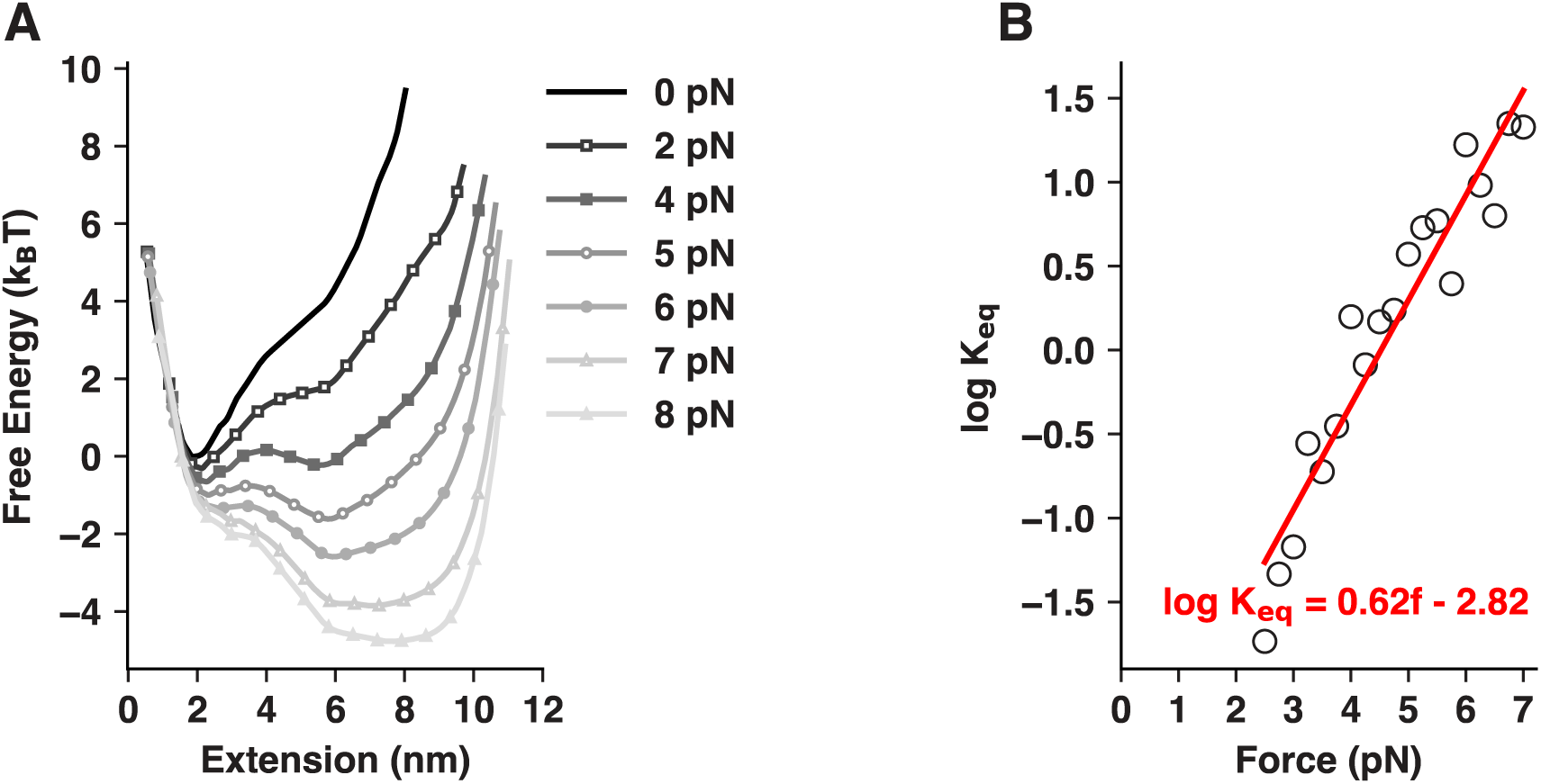
Simulated P5GA hairpin unfolding under constant pulling forces. (A) Free energy profiles as a function of end-to-end distance for P5GA hairpin at different pulling forces. (B) Logarithm of equilibrium constant as a function of pulling force, with a linear regression fit shown in red.

### Force-extension hysteresis of the ribozyme for estimating modeling timescales

Having established the accuracy of the SSPN model, we next examined the larger, experimentally characterized *Tetrahymena ribozyme*. A full-length version of this ribozyme has been studied using optical tweezers^46^ and an earlier CPU-based SSPN simulations.^42^ However, due to computational limitations, previous simulations required pulling speeds that were more than 3 orders of magnitude higher than those used experimentally, resulting in substantially higher rupture forces. Here, we revisit this system by focusing on its P5abc domain and calibrating the simulation speed to generate rupture forces comparable to the experimental values. We selected this subdomain of the full-length ribozyme because its thermodynamic and kinetic properties have been characterized in greater detail experimentally, including three variants that allow a complete analysis of the contribution of its constituent RNA motifs.^45^

To directly compare simulation with experiment, we modeled the three ribozyme variants – P5ab, P5abcΔA, and P5abc – that were studied in the optical tweezer experiment^45^ (Additional details of the modeling process are provided in Section *Modeling and simulations of P5ab, P5abc*ΔA*, and P5abc systems*). The mechanical unfolding of these three RNAs has been experimentally investigated,^45^ and these measurements provide the basis for validating and calibrating our SSPN simulations. Our simulation reveals a force-extension hysteresis when pulling apart two termini of the P5abcΔA RNA molecule (Figure 4A). The magnitude of this hysteresis allows us to estimate the timescale difference between the simulation and experiment. We calculated the simulated hysteresis range at different loading rates and compared it with the experimental measurement. As expected, hysteresis increases at a higher loading rate (Figure 4B), in agreement with the experimental observations.^45^ This analysis enables us to determine a reference loading rate *r*_ref_ = 5 × 10^8^ pN/s that reproduces experimental hysteresis measured at 10 pN/s. Hence, an estimated conversion factor of 5 × 10^7^ converts the simulation timescale to the experimental one.

**Figure 4:**
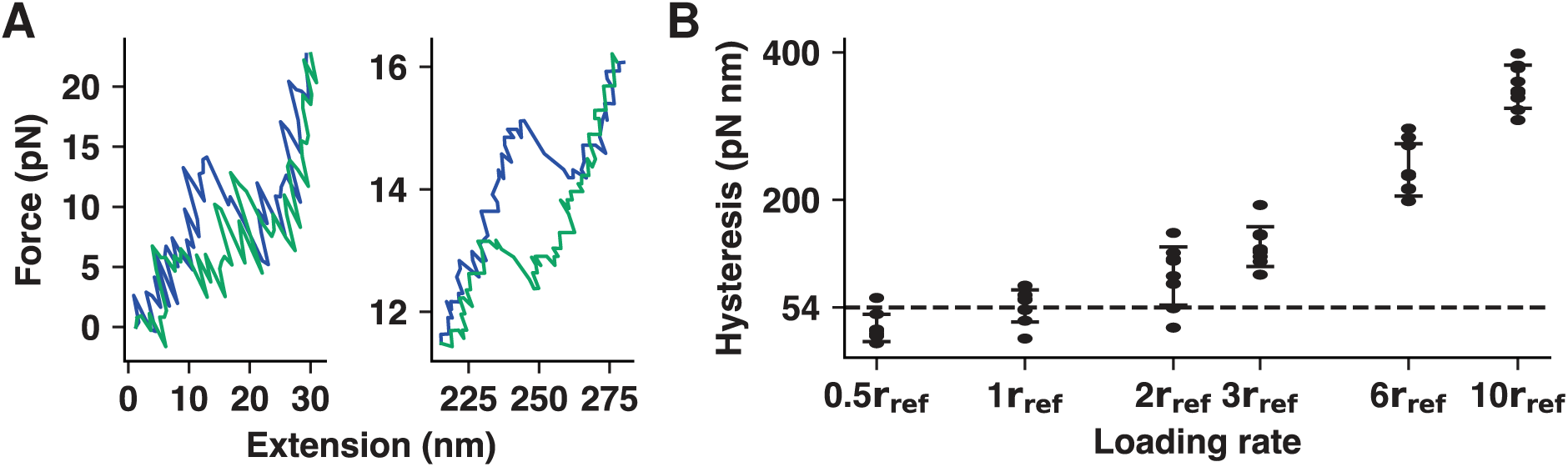
Time calibration of the SSPN model using P5abcΔA simulation. (A) Left: Simulated stretching (blue) and relaxing (green) force-extension curves. The hystere-sis is quantified as the area enclosed between the unfolding (blue) and refolding (green) trajectories. Right: Experimental force-extension curve reproduced from Reference 45 (the larger extension range comes from the long molecular handles in the optical tweezer).^45^ (B) Hysteresis values measured at six different loading rates, expressed relative to the reference loading rate *r*_ref_, which is defined as the rate at which simulated hysteresis matches the experimental value of 54 pN · nm (dashed line).

### SSPN model reproduced the unfolding free energies of ribozyme domain

Having calibrated the modeling time scales, we proceed to quantify the unfolding kinetics and thermodynamics of the P5abc systems. We first computed the equilibrium folding free energies of the three RNA ribozyme variants using umbrella sampling, ^62,64^ using Q^71^ – the fraction of native contacts formed – as the collective variable (Figure 5A, additional simulation details are provided in Section *Modeling and simulations of P5ab, P5abc*ΔA*, and P5abc systems*). Our simulations show that P5abc has the largest unfolding free energy penalty, followed by P5abcΔA and P5ab (Figure 5B). This trend is consistent with the fact that P5abc has the largest number of structure-based contacts introduced by the additional A-rich bulge and the P5c helix in P5abc. The quantified (un)folding free energy, Δ*G*_fold_ = 34 ± 7 *k*_B_*T*, lies within experimentally measured range (33-67 *k*_B_*T* ^45^). These results demonstrate that the SSPN model quantitatively captures RNA folding thermodynamics, consistent with previous studies using more detailed coarse-grained or all-atom structure-based models.^12,14,40^

**Figure 5:**
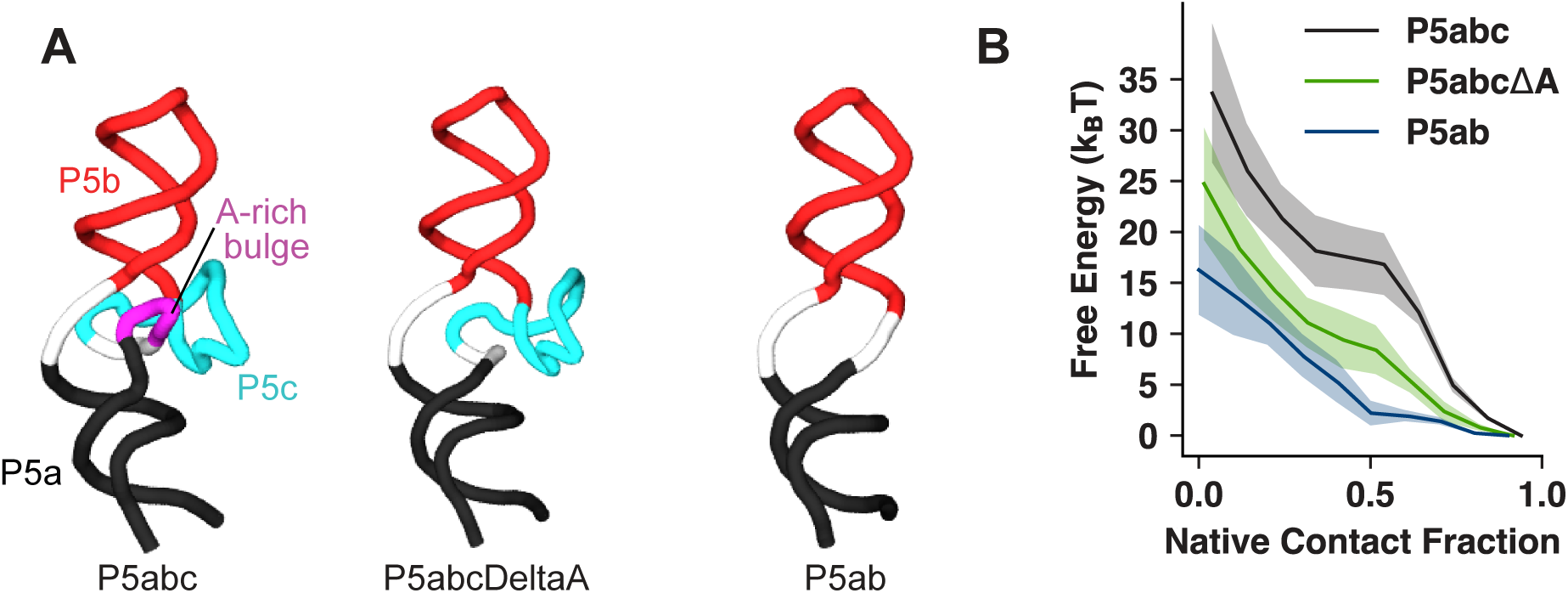
Structures of three P5abc variations and their unfolding free energy profiles. (A) Native structures of P5abc, P5abcΔA, and P5ab. (B) Free energy profiles of these systems as a function of Q, the fraction of formed native contacts. The corresponding free energies of unfolding are: 34±7 *k*_B_*T* for P5abc, 25±6 *k*_B_*T* for P5abcΔA, and 16±4 *k*_B_*T* for P5ab.

### SSPN reveals additional insights on the Folding pathways of P5abc ribozyme domain

Encouraged by the model’s ability to reproduce the thermodynamic properties of ribozyme domains, we next investigated the kinetic unfolding pathways of P5abc. We carried out simulations at a constant loading rate of *r*_ref_ = 5 × 10^8^pN/s, corresponding to 10 pN/s in the optical tweezer experiments.^45^ Our simulated rupture forces closely match experimental values (Figure 6A and B). Interestingly, while experiments suggest that rupture of the P5a helix is the rate-limiting step in cooperative unfolding,^45^ our simulations indicate that P5b and P5c helices are usually the kinetic bottleneck, whereas P5a unfolds at relatively lower forces (Figure 6A). To further characterize the kinetic behavior of unfolding P5abc, we computed the two-dimensional kinetic probability distribution of P5abc structural transitions from 200 constant-velocity simulation trajectories. These simulations reveal a dominant pathway in which P5a unfolds first, followed by P5b and P5c (Figures 6C and D). Unlike P5a, the order of P5b and P5c unfolding can be swapped, indicating two distinct kinetic pathways for P5abc to reach the fully unfolded random coil (Figure 6E). While topology plays a central role in determining the transition states in protein folding,^26^ the same may not apply to RNA, which is conformationally flexible.^11^ Therefore, accurate quantification of RNA unfolding pathways may require detailed characterizations of the sequence-specific energetics of RNA nucleotide interactions, thereby further improving our structure-based RNA model. Additionally, the presence of divalent Mg^2+^ ions is known to increase the P5a unfolding barrier.^45^ Explicitly modeling the effects of metal ions will likely correct the RNA unfolding order.

**Figure 6:**
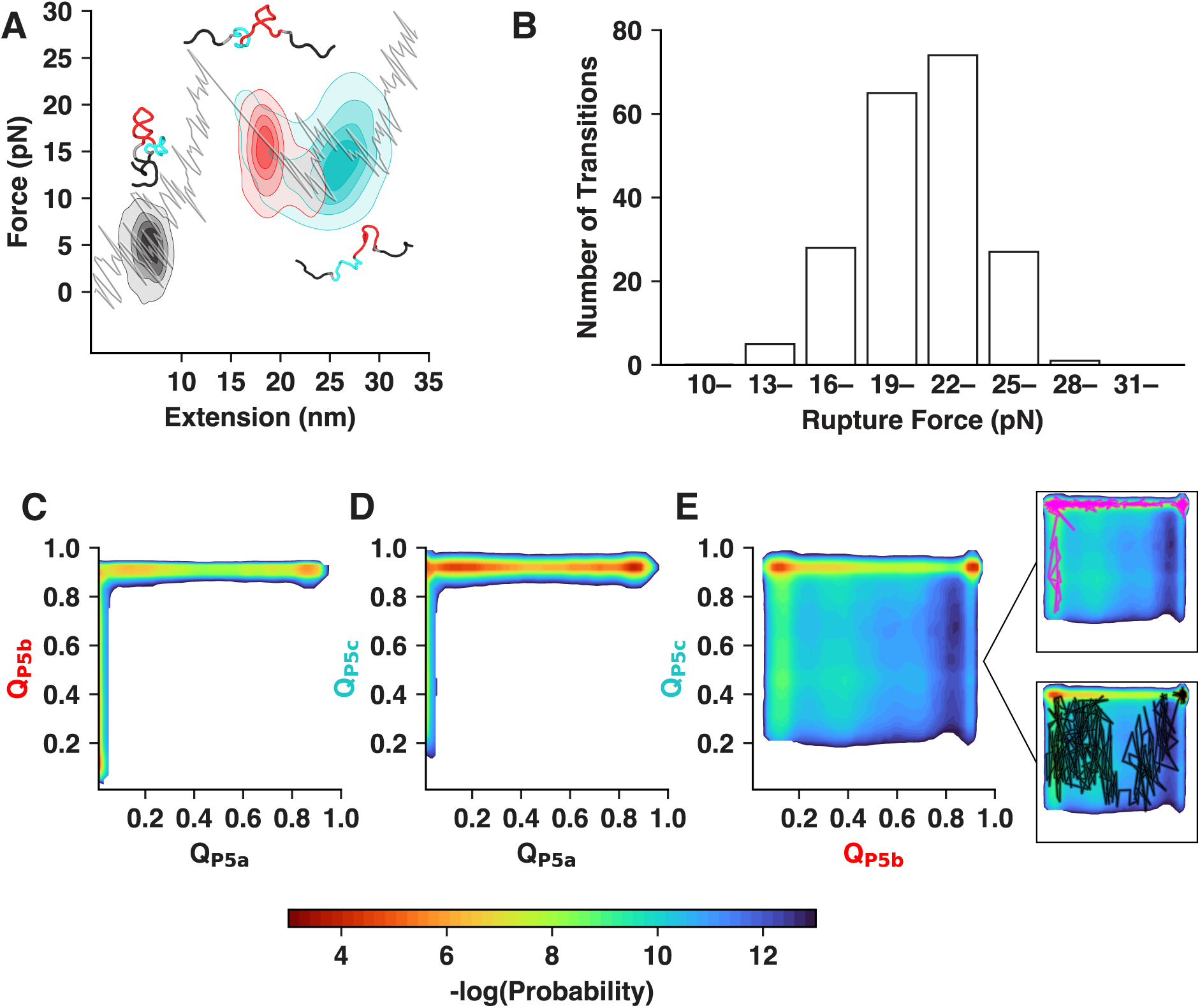
Kinetic unfolding pathways of P5abc. (A) Density of unfolding events for the three P5abc helices. A helix is considered unfolded when its Q*<*0.5. The contours denote the distribution of unfolding events. Representative structures at the moment of breaking each helix - P5a (black), P5b (red), and P5c (cyan) - are displayed, along with an example unfolding trajectory. (B) Statistics of rupture forces from 200 trajectories. The rupture force is defined as the force at the sharpest peak in the force-extension curve. (C-E) Two-dimensional kinetic probability distributions of structural transitions between fractions of native contacts formed in the P5a versus P5b helices (C), P5a versus P5c helices (D), and P5b versus P5c helices (E). Two representative trajectories (magenta and black) are highlighted in (E), illustrating two distinct unfolding pathways. All data were collected at the reference loading rate *r*_ref_.

## Discussion

In this manuscript, we implemented an SSPN structure-based model to examine RNA folding behavior. The model demonstrates high computational efficiency on a GPU-accelerated platform and can quantitatively capture both the folding thermodynamics and kinetics of RNA systems.

### Outlook for SSPN model to large RNA systems

The SSPN model achieves superior computational efficiency due to its coarse-grained and simplified force field without explicit solvent. Our computational benchmarks (Figure 2) show that the model can simulate over 1*µs* for systems containing around 400k of RNA nucleotide on a standard GPU, making it possible to simulate large RNA systems, including various long non-coding RNAs (lncRNAs)^6,7^ and collections of RNAs involved in LLPS,^9,47^ which typically contains of hundreds of RNAs with complex structures.

### Energetic details of the SSPN model

The SSPN model currently treats RNA-RNA interactions homogeneously based on RNA structures. This approximation has shown semi-quantitative accuracy in capturing RNA thermodynamics and kinetics, consistent with previous, more detailed, structure-based RNA models.^12,14,17,40^ Although this treatment ensures the model’s generalizability across various RNA folding systems, it may not fully capture the sequence-specific energetic details of RNA interactions. A future direction is to incorporate sequence-specific RNA-RNA interactions to better capture thermodynamic stability,^72^ and include sequence-specific protein-RNA interactions to improve modeling of protein-modulated RNA recognition and phase-separation behavior.^73–75^

### Extending SSPN to unstructured RNA

Many lncRNAs and phase-separating RNAs lack well-defined tertiary structures,^76,77^ posing a challenge for structure-based RNA models. To refine the SSPN model for simulating such RNAs, one can remove the tertiary structural native contacts while retaining backbone bonded interactions to preserve correct polymeric features. Similar strategies have been successfully applied to simulating disordered proteins, such as histone proteins.^51–53,57,78^ Careful calibration of nucleotide-nucleotide non-bonded interactions is essential to accurately characterize RNA dynamics.^43,79,80^ Additionally, experimental measurements, such as SHAPE^81^ and DMS^82^ data, or secondary structural predictions,^83–86^ can be used to constrain and regularize RNA secondary structure, thereby improving modeling accuracy.

### Explicit modeling of multivalent ions

Another area for future development is the incorporation of multivalent ions into the SSPN model, particularly Mg^2+^, which plays a central role in stabilizing tertiary structures and facilitates catalytic activities. ^24,87–90^ While the SSPN model currently accounts for Mg^2+^ implicitly through structure-based non-bonded potential, explicit modeling of multivalent ions is necessary to capture modulatory effects of these ions beyond Debye-Hückel treatment of charge screening by monovalent ions, including ion-ion correlations, ion multivalency, and entropic contribution by releasing ions.^54,55,91,92^ These effects may also underlie the experimentally observed high rupture forces and cooperativity of P5a helix unfolding,^45^ which are underestimated by the current model (Figure 6). Significant advances have been made in parameterizing explicit ion models for RNA simulations, ^24,25,93^ and these ion models can be incorporated into our SSPN model implementation.

## Acknowledgement

This work was supported by startup funding from North Carolina State University. Additional support was provided by the NC State Genetics and Genomics Academy and the Comparative Medicine Institute.

## Footnotes

∗ Changes in residue orientation can alter atomistic contacts. However, CG native pairs are defined by the existence of any atomistic native contacts between two residues and are therefore insensitive to modest residual reorientation.

## Notes

### Competing Interest Statement

The authors have declared no competing interest.

