## Supporting Information for "Efficient RNA Folding Simulation via a Structure-Based Single-Site-Per-Nucleotide Model"

#### Contents

**1 Figures** **S-2**

### 1 Figures

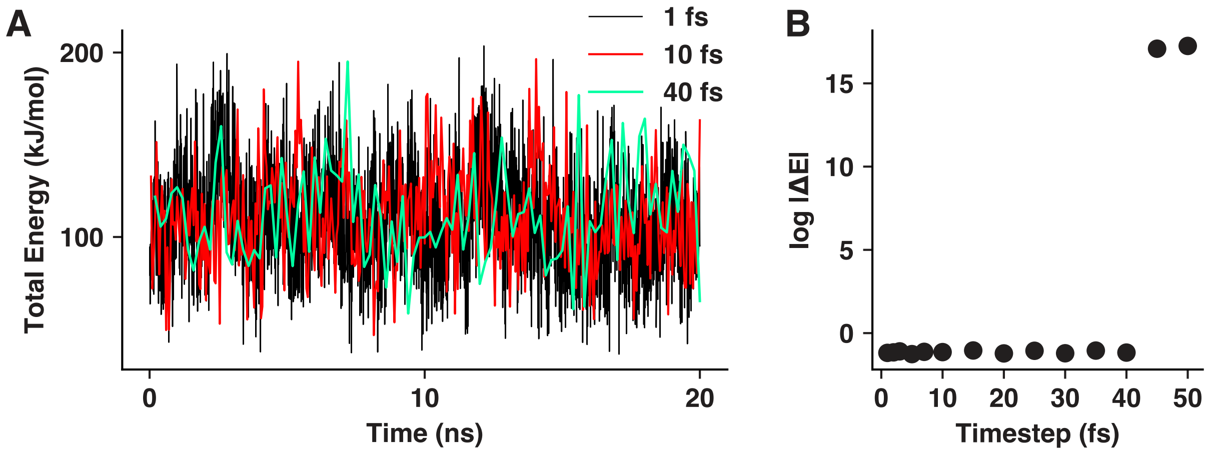


Figure S1: **Effect of timestep on simulation stability.** (A) Total energy of P5GA hairpin[S1,S2] as a function of simulation time across a range of timesteps. Only trajectories at 1 fs, 10 fs, and 40 fs were shown for clarity. (B) Energy drift as a function of timestep, where $|\Delta E| = \frac{1}{N}\sum_{k=1}^{N}|\frac{E_k - E_0}{E_0}|$, with $k$ being the frame index and $N$ being total number of frames.[S3] 40 fs is the largest timestep that maintains system stability, and we used 10 fs throughout all reported simulations.

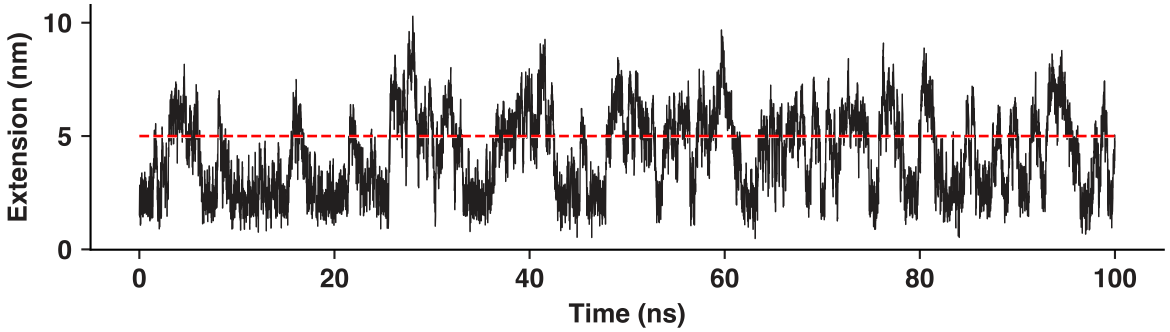


Figure S2: **End-to-end terminal residual distance of the P5GA hairpin as a function of time in constant force simulations at 4 pN.** The 5 nm threshold is highlighted in red, representing the unfolded state.

(S1) Rüdisser, S.; Tinoco, I. Solution structure of Cobalt(III)hexammine complexed to the GAAA tetraloop, and metal-ion binding to G.A mismatches. *J Mol Biol* **2000**, *295*, 1211–1223.

(S2) Hyeon, C.; Thirumalai, D. Mechanical unfolding of RNA hairpins. *Proc Natl Acad Sci U S A* **2005**, *102*, 6789–6794.

(S3) Knotts, T. A.; Rathore, N.; Schwartz, D. C.; De Pablo, J. J. A coarse grain model for DNA. *The Journal of Chemical Physics* **2007**, *126*, 084901.
